# Independent Co-option of a Tailed Bacteriophage into a Killing Complex in *Pseudomonas*

**DOI:** 10.1101/011486

**Authors:** Kevin L. Hockett, Tanya Renner, David A. Baltrus

## Abstract

Competition between microbes is widespread in nature, especially among those that are closely related. To combat competitors, bacteria have evolved numerous protein-based systems (bacteriocins) that kill strains closely related to the producer. In characterizing the bacteriocin complement and killing spectra for the model strain *P. syringae* B728a, we discovered its activity was not linked to any predicted bacteriocin, but is derived from a prophage. Instead of encoding an active prophage, this region encodes a bacteriophage-derived bacteriocin, termed an R-type syringacin. The R-type syringacin is striking in its convergence with the well-studied R-type pyocin of *P. aeruginosa* in both chromosomal synteny and molecular function. Genomic alignment, amino acid percent sequence identity, and phylogenetic inference all support a scenario where the R-type syringacin has been co-opted independently of the R-type pyocin. Moreover, the presence of this region is conserved among several other *Pseudomonas* species, and thus is likely important for intermicrobial interactions throughout this important genus.

## Importance

Evolutionary innovation is often achieved through modification of complexes or processes for alternate purposes, termed *co-option*. Similar co-options can occur independently in distinct lineages. Notable examples include the independent transition from forelimbs to wings in both birds and bats. Co-options can also occur at the molecular level within microorganisms. We discovered a genomic region in the plant pathogen *Pseudomonas syringae* that consists of a fragment of a bacteriophage genome. The fragment encodes only the tail of the bacteriophage, which is lethal towards strains of this species. This structure is similar to a previously described structure produced by the related species, *Pseudomonas aeruginosa*. The two structures, however, are not derived from the same evolutionary event. Thus, they represent independent bacteriophage co-option. The co-opted bacteriophage from *P. syringae* is found in the genomes of many other *Pseudomonas* species, suggesting ecological importance across this genus.

## Introduction

The genus *Pseudomonas* includes organisms exhibiting diverse ecological strategies and life histories, including human and plant pathogens, plant mutualists, as well as saprotrophic plant, soil, and water inhabiting organisms [1-6]. Many of the environments inhabited by this genus are limited in available forms of one or several key nutrients (e.g. carbon or nitrogen), which drives intermicrobial competition [7-9]. Furthermore, the distribution of nutrients can be spacially limited which drives competition for colonization of particular anatomical locations [10]. Resource competition is a major factor driving the evolution and ecology of microorganisms in many, if not all, environments.

Resource competition between microbes with overlapping niches can resolve in multiple ways, one of which relies on the production of anticompetitor compounds (i.e. compounds that directly inhibit ecological competitors) including bacteriocins [11,12]. Anticompetitor strategies are advantageous because they increase resource availability for the producer [13]. Bacteriocins are diverse, evolutionarily unrelated peptides, proteins, and protein complexes produced by bacteria that are lethal towards strains and species closely related to the producer [12,14]. As such, they are recognized as effective anticompetitor compounds. Indeed, numerous empirical and theoretical studies have demonstrated the adaptive advantage of bacteriocin production (e.g. [15-18]). In addition to basic scientific interest, there is considerable interest in harnessing and modifying bacteriocins as next generation pathogen control compounds [19-22]. Bacteriophage-derived bacteriocins (such as the R-and F-type pyocins) are of particular interest because of their ability to be reprogrammed to specifically target distantly related pathogens with few predicted side-effects [22].

Bacteriocins produced by *Pseudomonas aeruginosa* have been well documented (reviewed in [14,23]). Conversely, bacteriocins produced by environmental pseudomonads are less well understood, though this trend is changing (see [23] for a recent review). *Pseudomonas syringae* has served as a model species to understand many aspects of plant-microbe interactions, including the microbial ecology of aerial plant surfaces (termed the *phyllosphere*) as well as the molecular and genomic bases of pathogen-plant interactions [4,24-27]. Furthermore, there is a growing appreciation for the role of the water cycle in *P. syringae* dispersal and survival (reviewed in [28]), demonstrating the importance of environmental niches in the life cycle of this species. An important aspect of fitness in the leaf enviroment for *P. syringae* (and other plant-associated bacteria) is the ability to access and exploit favorable leaf anatomic locations (intercellular grooves, base of glandular trichomes) where water and nutrients are relatively abundant [24,29]. Such scarcity and distribution of resources drives competition among bacteria inhabiting the phyllosphere [7,30]. For certain coinhabitants, coexistence is mediated through resource partitioning, where dissimilarity between the catabolic potential allows the organisms to exploit distinct nutrient pools [7]. For coinhabitants with highly similar catabolic potential, however, resources are not effectively partitioned, and thus the microbes compete directly for the same niche leading to competitive exclusion. In these cases, as well as in similar situations where *P. syringae* likely competes for limited nutrients outside of the plant environment (such as in epilithic biofilms), we hypothesize that bacteriocins produced by this species will structure both evolutionary dynamics and community composition. Indeed bacteriocins have been predicted among the sequenced genomes of *P. syringae* (reviewed in [23, 31-34], with some members being biologically characterized [35].

In this work, we set out to assess the totality of bacteriocin content from 18 genomes representing the phylogenetic breadth of *P. syringae*, as well as assess the killing activity spectra from these same strains. In attempting to link the genotype to phenotype in the model strain, *P. syringae* B728a, we discovered its killing activity could not be explained by any predicted bacteriocins. In searching for the source of this killing activity, we have discovered a new bacteriophage-derived bacteriocin, or tailocin (as per the terminology used in [23,36,37]). This killing compound represents striking genomic and molecular convergence with the R-type pyocin of *P. aeruginosa*, but is independently derived. Additionally, we show that this tailocin is largely conserved throughout *P. syringae*, and is present in other related *Pseudomonas* species, though its genomic location is variable. Thus, this killing compound is likely a major mediator of intraspecific interaction across this ecologically important genus.

## Results

### P. syringae strains commonly exhibit killing activity and harbor diverse predicted bacteriocin loci

Mitomycin C induction resulted in production of diverse killing patterns from *P. syringae* pvs. in culture supernatants, as evidenced by a zones of inhibition created by filtered supernatants in a lawn of target strains. The killing activity could be the result of a bactericidal compound or an activated prophage, which are commonly predicted in sequenced *P. syringae* genomes and have been recovered using this protocol (e.g. [38,39]). Only two strains produced detectable bacteriophage activity (Fig. S1). Though we observed abundant bacteriocin-mediated clearing, we can’t rule out the possibility that bacteriophages were more prevelant, but were not detected because their activity was masked by that of a bacteriocin(s) or that induced bacteriophage could not infect strains included in this study.

The genomes of all strains were analyzed using several approaches (see methods). Class III bacteriocins (> 10 kDa) with DNase catalytic domains (colicin E9-like and carocin D-like) were commonly predicted among the strains (data not shown). Additional class III bacteriocins predicted among strains include pyocin S3-like (DNase), colicin E3-like (rRNase), colicin D-like (tRNase), colicin M-like (lipid II degrading), and putadicin-like (unknown killing mechanism). In addition to class III bacteriocins, there were several class I and II, collectively referred to as ribosomally-synthesized and post-translationally-modified peptides (RiPPs), including microcin-like, lasso peptide-like, sactipeptide-like, and linear azol(in)e-containing peptide-like peptides and associated modification and transport proteins. The BAGEL2 and BAGEL3 databases used for bacteriocin prediction contain criteria for all major bacteriocin classes except for bacteriophage-derived bacteriocins, such as the R-and F-type pyocins produced by *Pseudomonas aeruginosa*[40,41].

We sought to confirm the genetic basis of the killing activity produced by *Psy* B728a. To this end, we constructed targeted gene deletions for the two predicted colicin-like bacteriocins, Psyr_0310 and Psyr_4651 (hereafter referred to as s-type syringacins after the naming convention established for *P. syringae* and *P. aeruginosa*[14,35]). Both s-type syringacins encode putatively DNase-active proteins. Deletion of Psyr_0310 and Psyr_4651, individually or in tandem, resulted in no detectable loss of killing activity against any strain targeted by the wild type *Psy* B728a (Fig. 1). This result indicated that the killing activity produced by *P. syringae* B728a is derived from a source not predicted by genome mining nor blastp queries with conserved catalytic domains.

**Figure 1.**
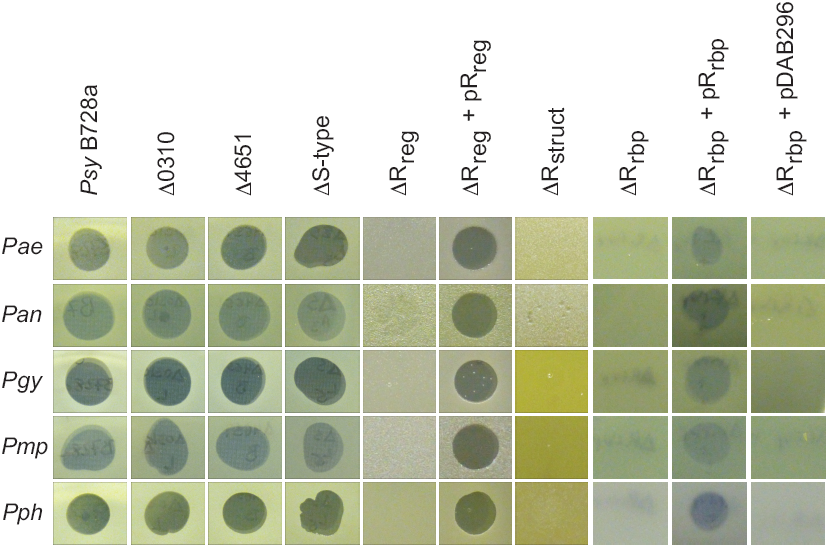
Clearing activity produced by *Psy* B728a and various bacteriocin deletion mutants against select pathovars. 2-5 µl of filter-sterilized, mitC induced culture supernatant was spotted onto soft agar overlays of the indicated pathovars (rows). *Psy* B728a (wild type); Δ0310 (deletion of Psyr_0310); Δ4651 (deletion of Psyr_4651); ΔS-type (deletion of both Psyr_0310 and Psyr_4651); ΔR_reg_ (deletion of Psyr_4603 promoter and coding sequence, deletion of Psyr_4602 promoter); ΔR_reg_ + pR_reg_ (R_reg_ deletion with both Psyr_4603 and Psyr_4602, as well as surrounding genomic sequence, complemented *in trans* on a replicative vector); ΔR_struct._ (deletion of Psyr_4592 through Psyr_4586, as well as the promoter of Psyr_4585); ΔR_rbp_ (deletion of Psyr_4584 and Psyr_4585); ΔR_rbp_ + pR_rbp_ (Psyr_4584 and Psyr_4585 expressed from a constitutive promoter in pBAV226); ΔR_rbp_ + pDAB296 (pBAV266 empty vector).

### P. syringae B728a harbors a prophage-like region, which is located in a syntenic region with the R and F-type pyocins encoded by Pseudomonas aeruginosa, that is responsible for its killing activity

The above results indicated that another locus (or loci) was responsible for the observed killing activity from *Psy* B728a. As the killing activity produced by this strain was more abundant from cultures treated with mitomycin C, we looked for the presence of RecA-mediated autopeptidase regulators, which are known to regulate bacteriocin production in *P. aeruginosa* and are mitomycin C responsive [42]. There are 9 open reading frames predicted in the Psy B728a genome that exhibit significant similarity to the consensus sequence for COG1974 (SOS-response transcriptional repressors) retrieved from the NCBI Conserved Domain Database (CDD) [43]. One of the predicted RecA-mediated autopeptidases was found associated with the previously described prophage region II [44], which is syntenic to that of the R-and F-type pyocins (specifically between *trpE* and *trpG*) of *P. aeruginosa* PAO1 (Fig. 2). As both the R-and F-type pyocins are derived from bacteriophages [45], we hypothesized that prophage region II was responsible for the mitomycin C inducible killing activity. We constructed a series of deletion in both structural and regulatory genes associated with prophage region II (Fig. S2 A and C). Deletion of a putative transcriptional regulator (Psyr_4603), encoding a RecA-mediated autopeptidase, as well as the promoter of a divergently transcribed hypothetical protein (Psyr_4602) resulted in loss of killing activity against all strains sensitive to *Psy* B728a (Fig. 3). Because activity of both Psyr_4603 and Psyr_4602 are affected in this deletion strain, we refer to it as ΔR_reg_ (indicating the potential deletion of multiple regulatory gene functions) Complementation by introduction of Psyr_4603 and Psyr_4602 (as well as flanking genomic sequence) in trans resulted in restored killing activity against all strains tested. Deletion of a region encompassing Psyr_4585 through Psyr_4592, which includes genes encoding structural components of baseplate, spike, and tail fiber, and tape measure (ΔR_struct_), resulted in abolished killing activity against all strains. Deletion of the predicted receptor binding protein (Psyr_4585) and chaperone (Psyr_4584) that assists in attachment of the receptor binding protein to the baseplate (ΔR_rbp_), also resulted in abolished killing activity against all strains. Expression of Psyr_4585/Psyr_4584 from a constitutive promoter in trans resulted in restored killing activity against all strains. It should be noted that complementation was only possible for constructs which included an upstream alternative start codon (Fig. S2B). Additionally, electron microscopy showed the presence of bacteriophage tails lacking capsids in culture supernatants of the wild type strain but not in the supernatants of either ΔR_reg_ or ΔR_struct_ (Fig. S2D). Taken together, the genetic and phenotypic data convincingly demonstrate the bacteriophage-derived region is responsible for the intraspecific killing activity produced by *Psy* B728a. In keeping with nomenclature for *P. aeruginosa*, we refer to the killing complex produced by *P. syringae* as an R-type syringacin.

**Figure 2.**
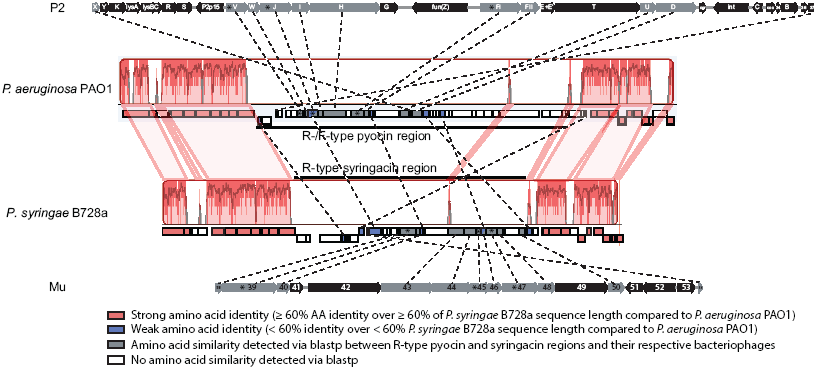
Comparison of genomic regions flanked by *trpE* and *trpG* between *P. syringae* B728a and *P. aeruginosa* PAO1. Regions between *P. aeruginosa* PAO1 and *P. syringae* B728a connected pink shading indicate regions of significant nucleotide identity as assessed by progressive Mauve. Dashed lines were omitted between conserved genes in regions of nucleotide identity between *P. aeruginosa* PAO1 and *P. syringae* B728a, however gene synteny between the two strains is conserved. Dashed lines between bacterial and bacteriophage genomes indicate genes exhibiting amino acid similarity detectable by blastp. Genes marked by asterisks indicate those utilized for the phylogenetic analyses.

**Figure 3.**
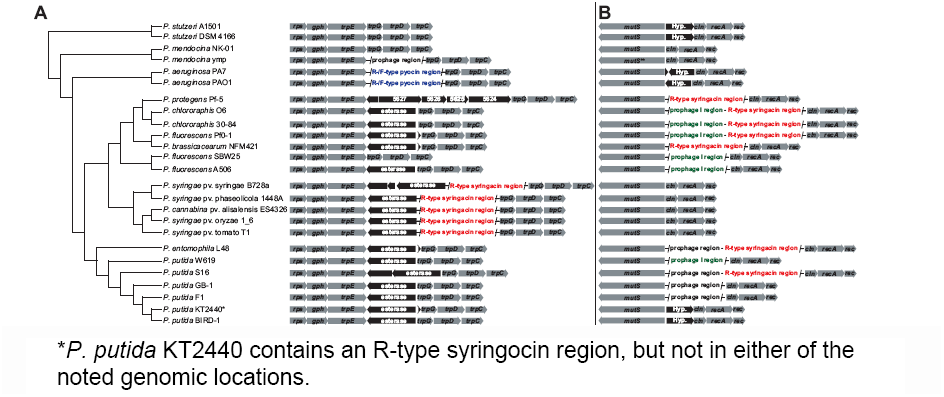
Comparison of gene content between *trpE* and *trpG* (A) or between *mutS* and *cinA* (B) across select *Pseudomonas* species. Phylogeny is adapted from **[46]**which was constructed using maximum likelihood based 726 proteins shared across the genomes. Genes and genomic regions are colored as follows: conserved genes demarcating the genomic location (gray arrows), non-phage related genes (black arrows), R-type syringacin-like region (red), R-and F-type pyocin-like regions (blue), prophage I region described in **[46]**(green), undescribed prophage regions (black). * *P. putida* KT2440 contains an R-type syringocin region, but not in either of the noted genomic locations.

### The R-type syringacin of P. syringae is co-opted from an independent bacteriophage, and thus is evolutionarily convergent with the R-type pyocin at genomic, molecular, and phenotypic levels

Despite the conserved synteny between the R-type pyocin and R-type syringacin, few genes in these regions exhibit similarity detectable using a blastp search with low stringency (E-value cut off ≤10^-5^) (Fig. 2). The R-type syringacin of *Psy* B728a was originally described as being related to a Mu-like bacteriophage based on gene content and arrangement [44]. To further confirm that the R-type syringacin is not derived from the same progenitor of the R-type pyocin of *P. aeruginosa*, we compared the gene content between *trpG* and *trpE* across multiple pseudomonads (Fig. 3A). Across the *Pseudomonas* phylogeny (as described in [46]), the gene content between *trpE* and *trpG* is variable. Importantly, the R-type syringacin region located between *trpE* and *trpG* is only found in the *P. syringae* clade; the other pseudomonads closely related to *P. syringae* including *Pseudomonas putida*, *Pseudomonas flurosescens*, and *Pseudomonas chlororaphis* do not harbor any predicted prophage or prophage-like elements in this region. Interestingly, however, many of the strains related to *P. syringae* do harbor an R-type syringacin region, but at an alternate genomic location (Fig. 3B, see below).

Additionally, a nucleotide alignment of the sequences surrounding and including the R-type pyocin and syringacin regions demonstrated high nucleotide sequence identity between the two strains for *trpE* and *trpG* (including genes operonic with these loci), but negligible nucleotide identity corresponding to the regions encoding the respective R-type bacteriocins (Fig. 2). This result corresponds well with the poor amino acid similarity between the ORFs of these regions: those genes located in the *trpE* and *trpG* operons are well conserved at the amino acid level between the two strains, whereas very few genes within the respective bacteriocin regions have detectable amino acid sequence similarity, and those that do exhibit relatively little similarity (Fig. 2).

To further confirm that the R-type syringacin from *P. syringae* is derived independently of the R-type pyocin from *P. aeruginosa*, we performed phylogenetic reconstructions using a maximum likelihood-based (ML) approach for three proteins where amino acid similarity could be detected by blastp across *P. syringae* B728a, *P. aeruginosa* PAO1, phage Mu, and phage P2 using a transitive homology approach [47]. Based on the functions predicted or described for the orthologs from P2 and Mu [48,49], Psyr_4587 encodes a component of the baseplate, Psyr_4589 encodes the spike, and Psyr_4595 encodes the tail sheath protein. For all proteins, there was overwhelming support that the orthologs from *P. syringae* B728a (as well as all other *P. syringae* strains examined in this study, except for those where the protein was truncated [see below]) were more closely related to those from phage Mu (with 96-100 bootstrap support (bs)), whereas the orthologs from *P. aeruginosa* were more closely related to those from phage P2 (92-100 bs; Figs. 4, S3, and S4). Thus, the phylogenetic inference supports that R-type syringacin is derived from an independent bacteriophage (a Mu-like bacteriophage) rather than the R-type pyocin, despite both R-type bacteriocins being located in a syntenic genomic region. These results are similar to those reported previously, where Mavrodi et al. found the R-type syringacin region (therein reported as genomic island 12) from *P. syringae* B278a exhibited nucleotide similarity to prophage 1 of *P. protegens* (*P. fluorescens*) Pf-5, whereas the R-type pyocin of *P. aerugionas* PAO1 exhibited nucleotide similarity to prophage 3 of Pf-5 [50].

**Figure 4.**
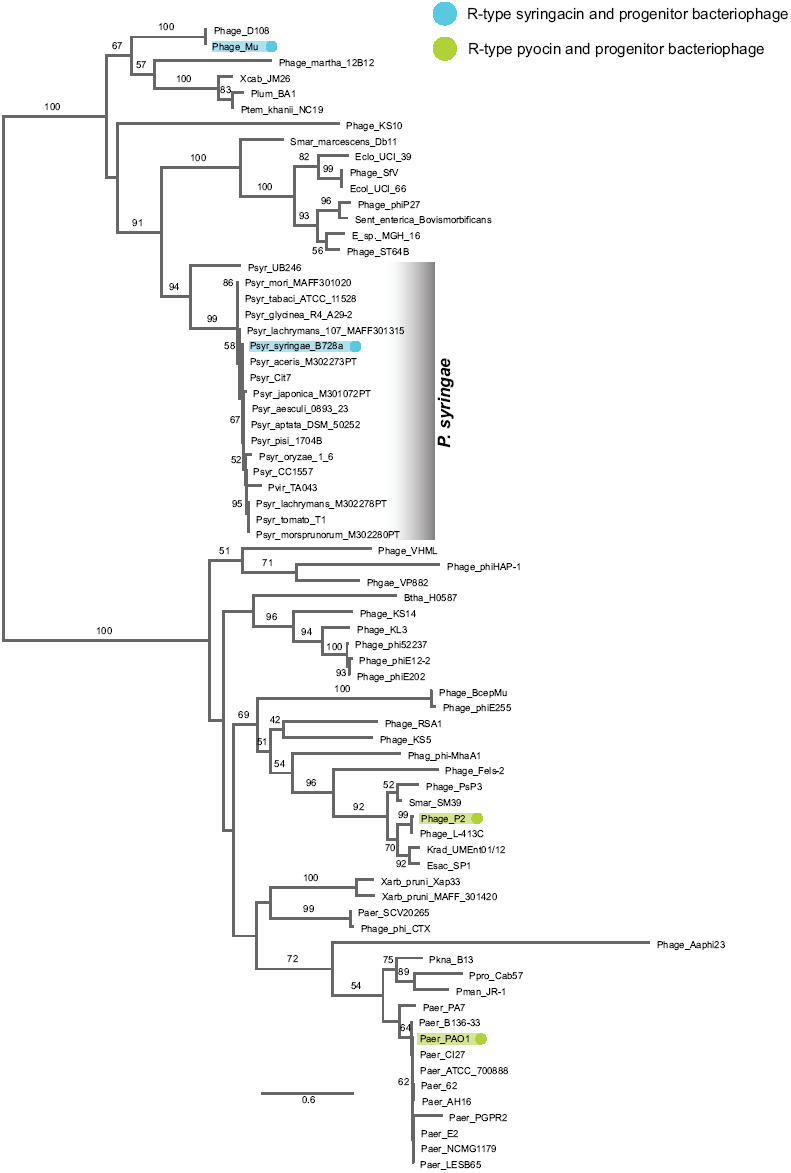
Maximum likelihood phylogeny of Psyr_4589 (*P. syringae* B728a), PA0616 (*P. aeruginosa* PAO1), gp45 (Mu), and gpV (P2) and related protein sequences from bacteria and bacteriophages. Teal highlighted nodes indicate Psyr_4589 and gp45, whereas green highlighted nodes indicate PA0616 and gpV. Values on branches indicate bootstrap support (1000 bootstrap replicates) for those clades (only values of 50 or greater are shown). A monophyletic clade including all *P. syringae* orthologs is indicated. See table S2 for accessions or IMG GeneIDs associated with each sequence.

Though our analyses indicate the R-type syringacin has been derived independently of the R-type pyocin, comparing the gene content of the R-type syringacin to phage Mu demonstrates the loss and retention of similar genes to that described for the R-type pyocin. Specifically, the R-type syringacin lacks all genes corresponding to the early and middle region of the phage Mu genome (encoding functions related to phage replication). The genes retained that correspond to the late region of phage Mu, are only those related to tail morphogenesis, whereas late genes related to capsid morphogenesis and DNA packaging have been lost (Fig. 5). This pattern of gene retention mirrors that of the R-type pyocin when compared to phage P2 [45]. These results indicate convergent bacteriophage co-option at both genomic and phenotypic levels.

**Figure 5.**
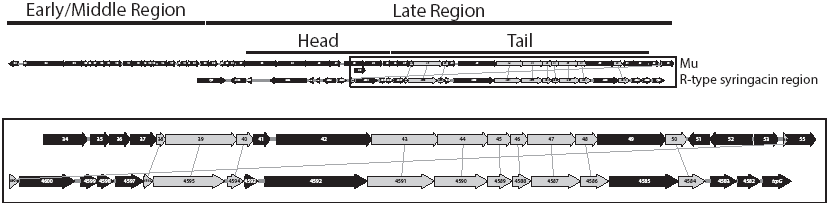
Comparison of gene content in R-type syringocin region and bacteriophage Mu genome. Genes exhibiting detectable amino acid similarity by blastp are indidcated in gray. Designation of phage Mu gene categories (early, middle, late, head, tail) taken from **[48]**.

Though we have compared the R-type syringacin to the tail of bacteriophage Mu, it appears that it is most closely related to the tail of serotype converting phage SfV of *Shigella flexneri*, based on phylogenetic inference and overall results from blastp comparisons (i.e. there is a greater number of hits to the SfV tail, as well as greater amino acid identity for hits to SfV than to Mu) (Figs 4, S3-S4, and table S1). Importantly, phage SfV harbors a Mu-like tail [51]. In contrast to genes encoding components of the phage tail, prediction of genes associated with lysis functions was more complicated. No gene within this region displays sequence similarity to any described holin. There are two predicted chitinases (Psyr_4583 and Psyr_4600) within this region, which may harbor peptidoglycan-degrading (endolysin) activity as chitinases have been shown to exhibit such activity [52,53], and chitinase domains have been predicted within phage genomes previously [54,55]. We were able to predict one spannin-encoding gene (Psyr_4582). Thus, there is currently an incomplete accounting of all required lysis functions within this region. This may result from lysis functions being provided by genes within this region that lack sequence similariy to known lysis genes, or lysis functions may be provided in trans from prophage region I, which harbors an active prophage (Hockett and Baltrus, unpublished results).

### The R-type syringacin is conserved across Pseudomonas

Although the gene content between *trpE* and *trpG* across the *Pseudomonas* strains presented in figure 3A indicated that only *P. syringae* and *P. aeruginosa* harbor R-type bacteriocins, a previous report [50] demonstrated nucleotide conservation between prophage 1 of *P. fluorescens* Pf-5 (recently renamed to *P. protegens* Pf-5, [56]) and the R-type syringacin (referred to as genomic island 12). We confirmed the homology of these regions by comparing the amino acid conservation and synteny of genes in their respective regions (data not shown). There is significant similarity at the amino acid level between most of the tail structural genes. Additionally, the order of genes is syntenic between the two organisms. The most dissimilar proteins between the two regions are those encoding the receptor binding proteins (also referred to as the tail fibers), Psyr_4585 and PFL_1226, exhibiting only 66% amino acid similarity over 8.6% of the length of Psyr_4585). This result is not unexpected, however, given this protein's role in host recognition, and that it is a highly divergent protein between related bacteriophages as well as between R-type pyocins encoded among strains of *P. aeruginosa*[23,57]. Our results in conjunction with those previously reported, indicate that prophage I of *P. protegens* Pf-5 is homologous to the R-type syringacin, and thus likely encodes a killing compound within this organism.

As previously shown [46,50] prophage I of *P. protegens* Pf-5 is similar to prophage regions found in closely related *P. fluorescens* strains as well as other pseudomonads. These previous results indicate that R-type syringacin-like elements exist in *P. chlororaphis* 30-84, *P. chlororaphis* O6, and *P. fluorsecens* PfO-1. Inspecting the genomes of all strains shown in figure 3 using blastp with several R-type syringacin genes used as the queries, further demonstrated R-type syringacin regions present in *P. brassicacearum* NFM421, *P. entimophila* L48, *P. putida* S16, and *P. putida* KT2440 (Fig. 3B).

Phylogenetic analysis of this region, using concatenated homologs of Psyr_4587, Psyr_4589, and Psyr_4595, across these strains indicates largely, but not entirely, vertical inheritance (Fig. 6). Clustering of *P. entomophila* with *P. fluorescens*, *P. brassicacearum*, and *P. chlororaphis*, instead of with the *P. putida* strains, to which *P. entomophila* is more closely related, for all individual protein phylogenies (Fig. S5) as well as the concatenated phylogeny (Fig. 6) is well supported (≥ 70 bs). This suggests that the R-type syringacin was co-opted by *P. entomophila* from the ancestor of the *P. fluorescens* clade. The placement of *P. syringae* UB246 is incongruent among the phylogenetic reconstructions. *P. syringae* UB246 is an early diverging strain of this species [6,58]. The concatenated phylogeny (with poor bs) as well as the phylogeny of Psyr_4595 orthologs (with strong bs) indicate the *P. syringae* UB246 R-type syringacin is more closely related to the R-type regions from *P. putida*, than it is to the R-type regions of the other *P. syringae* strains. The single gene phylogenies for Psyr_4587 and Psyr_4589 place the *P. syringae* UB246 ortholog on a branch including, and sister to, all of the strains except *P. putida* strains, or outside of all the *Pseudomonas* species, including *P. putida*, respectively. Thus, the exact relationship of the R-type syringacin of *P. syringae* UB246 to the other R-type regions is currently ambiguous. Comparison of this region across the *P. syringae* strains phenotypically characterized in this study revealed broad conservation (Fig. 7).

**Figure 6.**
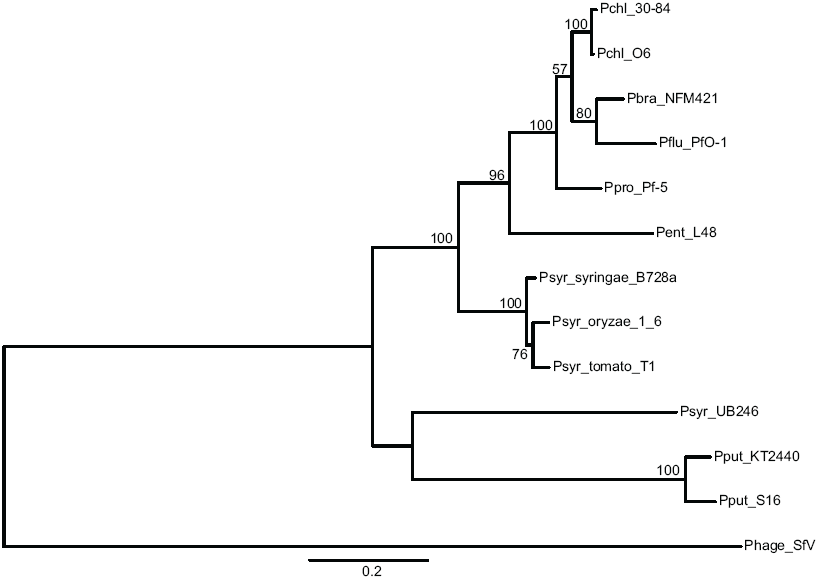
Maximum likelihood phylogeny of concatenated amino acid sequences of Psyr_4587, Psyr_4589, and Psyr_4595, and their homologs from *Pseudomonas* strains harboring an R-type syringocin-like region. Values indicate bootstrap support (out of 1000 replicates, only values greater than 50 are shown). The following strains are Pchl 30-84 (*P. chlororaphis* 30-84), Pchl O6 (*P. chlororaphis* O6), Pbra NFM421 (*P. brasicacearum* NFM421), Pflu PfO-1 (*P. fluorescens* PfO-1), Ppro Pf-5 (*P. protegens* Pf-5), Pent L48 (*P. entomophila* L48), Psyr B728a (*P. syringae* pv. syringae B728a), Psyr 1_6 (*P. syringae* pv. oryzae 1_6), Psyr T1 (*P. syringae* pv. tomato T1), Psyr UB246 (*P. syringae* UB246), Pput KT2440 (*P. putida* KT2440), Pput S16 (*P. putida* S16). Phage SfV (bacteriophage SfV, outgroup).

**Figure 7.**
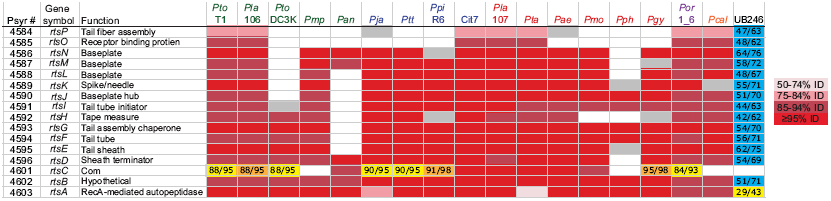
Conservation of R-type syringacin across *P. syringae*. Cells colored in shades of red indicate gene presence with the noted % amino acid identity to the *Psy* B728a homolog, which are syntenic to the R-type syringacin in *Psy* B728a. Cells noted in gray are syntenic to the R-type syringacin of *Psy* B728a and exhibit amino acid similarity detectable by blastp, but have less than 60% query coverage (potentially indicating the gene is truncated compared to the ortholog in *Psy* B728a). Yellow cells indicate genes that exhibit significant amino acid identity and similarity (%identity/%similarity) to the noted gene in *Psy* B728a, however the genes are present in a non-syntenic region of the noted genome. Orange cells indicate genes that exhibit significant amino acid identity and similarity to the noted gene in *Psy* B728a, however their location within their respective genomes can‘t be determined. Blue cells (UB246) indicate genes that exhibit detectable similarity those indicated in *Psy* B728a, additionally the genes are present in the same order as those in the R-type syringacin, however, they are located in a different genomic region (not flanked by *trpE* and *trpG*). Functional assignments are based either on amino acid sequence similarity either to phage Mu or phage SfV (*rtsC*-*rtsN*, *rtsP*), synteny with described genes from either phage (*rtsO*), or gene annotation at Integrated Microbial Genome website (*rtsA*, *rtsB*).

### Naming of genes associated with the R-type syringacin

As we have described the function and a portion of the molecular determinants for a bacteriophage derived bacteriocin in *P. syringae* B728a, we have named genes associated with this complex to *rtsA*-*J* for <u>R</u>-<u>t</u>ype <u>s</u>yringacin (see Fig. 7). Our aim in naming these genes is two-fold. First, providing uniform nomenclature for this structure reflects its specialization as a killing complex, and not simply a degenerate prophage. Additionally, we hope this nomenclature will aid researchers in distinguishing this region from the R-type pyocin, especially when discussing homologous tailocins produced by non-*syringae* and non-*aeruginosa* pseudomonads (e.g. *P. fluorescens*, *P. putida*).

## Discussion

Given the diversity and distribution of bacteriocins across *P. syringae*, as well as related pseudomonads [23,34,46], it is highly likely that these systems mediate ecological and evolutionary interactions between organisms in the plant environment and beyond. Indeed, that Mavrodi *et al.* (2002, 2009) found the region harboring the R-type bacteriocin present in *P. fluorescens* strain Q8r1-96 but absent in strain Q2-87, which is less competent at colonizing the wheat rhizosphere, indicates that this region could affect root colonization [50,59]. Additionally, Garrett et al. found a general trend where *P. syringae* isolates recovered from citrus tended to produce bacteriocins active against isolates recovered from pear and vice versa [60]. Direct demonstration of the benefit of bacteriocins in plant colonization is still lacking, and thus is an area ripe for future investigation.

Previous work by Fischer et al. demonstrated the presence of a bacteriophage tail-like bacteriocin produced by *P. fluorescens* SF4c, which killed related *P. fluorescens* strains [61]. Comparing the results of Fischer et al. with Loper et al., the phage-tail bacteriocin of SF4c is related to the prophage I in Pf0-1, as described in [46]. In Pf0-1, this region is located immediately upstream of prophage II, which is related to the R-type syringacin described in this study. Thus, it appears that tailocins are more common and widespread throughout *Pseudomonas* than previously recognized. Indeed, previous studies into bacteriocins of *P. syringae* strains indicated the production of high molecular weight bacteriocins (synonymous with tailocins) [62], including reports of phage-like particles, such as syringacin 4-A [39]. To our knowledge, however, research from this era did not characterize the genetic basis of these proteins. That our genomic predictions initially only identified low molecular weight bacteriocins, but not the R-type syringacin, likely reflects the relative difficulty in predicting the latter compared to the former. Indeed, despite the R-type syringacin and R-type pyocin being functionally similar and located in the same region in their respective genomes, that they display negligible gene nucleotide or amino acid identity would preclude prediction using sequence similarity. Thus, *de novo* prediction of tailocins from genomic sequence is a difficult task, unless the tailocin is closely related to one that has been previously characterized.

Both this and previous work found R-type syringacin-like regions throughout the *Pseudomonas* intragenic cluster II [63], including *P. protegens*, *P. chlororaphis*, *P. fluorescens*, *P. brassicacearum*, *P. putida*, *P. entomophila*, and *P. syringae*[46,50]. The R-type syringacin is widely distributed through out this group of related pseudomonads with a complex evolutionary history. This region appears after the divergence of *P. aeruginosa* from other species. In *P. syringae*, the R-type region is found nearly exclusively between *trpE* and *trpG*, the only exception being strain UB246, an early diverging member of *P. syringae sensu lato*[58], indicating a single introduction into this species. Supporting the conclusion of a single introduction into *P. syringae*, in all phylogenies, the UB246 orthologs are sister to all other *P. syringae* proteins, likely indicating the introduction occurred prior to the diversification of this species. A parsimonious explanation for the distribution of the tailocin within the *P. fluorescens*/*P. chlororaphis*/*P. protegens* clade is that it was ancestral, being subsequently lost by *P. fluorescens* strains SBW25 and A506, which constitute a monphyletic clade (with 100 bootstrap support as reported in [46], Fig. 3). The pattern of retention in *P. putida*/*P. entomophila*, however, is more complex. Several strains of *P. putida* do not contain the R-type syringacin region (strains W619, GB-1, F1, BIRD-1), whereas strains S16 and KT2440 do contain this region, though in different genomic locations. Additionally, despite *P. entomophila* being more closely related to *P. putida*, all phylogenetic analyses supported its R-type region being more closely related to that from *P. fluorescens*/*P. chloroaphis*/*P. protegens*. Thus, in addition to the intragenomic mobility of this region, there appears to be intergenomic mobility as well.

The results presented above clearly demonstrate the co-option of a retractile-type bacteriophage into a retractile-type bacteriocin independent of the R-type pyocin of *Pseudomonas aeruginosa*. At the DNA level, there is a high degree of identity between *P. aeruginosa* and *P. syringae* in the *trpE* and *trpG* (and surrounding genomic regions) that is absent within the corresponding bacteriocin regions. Additionally, amino acid sequence similarity between the genes associated with each bacteriocin is scant, and likely reflects similarity based on conserved function, as both bacteriocins are derived from related bacteriophages (both in the *myoviridae* family) [49]. Finally, the phylogenetic reconstruction based on three structural proteins, all provided strong support for separating all of the *P. syringae* R-type syringacin sequences into a monophyletic clade with bacteriophage Mu, whereas all of *P. aeruginosa* R-type pyocin sequences were present in a monophyletic clade with bacteriophage P2, thus indicating they are independently derived.

The convergence between these two bacteriocins is striking at both the genomic and molecular levels. Given that the R-type syringacin region is located in at least 3 (possibly 4) distinct genomic locations across all of the strains included in this study, this region has likely mobilized in certain lineages following co-option. Currently, however, we cannot conclusively determine which location was the ancestral location where the prophage was originally co-opted to a tailocin. The distinct locations include between *trpE* and *trpG* (*P. syringae* pathovars), between *mutS* and *cinA* (*P. fluorescens* and closely related strains, *P. entomophila*, *P. putida* S16), and in currently undefined regions in *P. syringae* UB246 and *P. putida* KT2440. It is particularly interesting that the tailocin of *P. syringae* pathovars and the tailocin of *P. aeruginosa* are found in syntenic genomic locations, despite being derived independently. It is currently unclear what features favor mobilization or integration into this region. In any event, the convergence of these events implies there is selective pressure for genetic integration at this locus in *Pseudomonas*. At the molecular level, there is a similar profile of gene retention and loss between the R-type pyocin and R-type syringacin. Gene encoding functions related to replication, capsid morphogenesis, and DNA packaging have been lost in the tailocins, with only genes encoding functions related to the bacteriophage tail morphogenesis being retained. Again these results point to a strong selective pressure favoring the co-option of bacteriophages into killing complexes. This work contributes to the body of research demonstrating the adaptive potential provided by bacteriophage to their bacterial hosts [37,64].

## Materials and Methods

### Plasmids, primers, bacterial isolates, and growth conditions

All bacterial strains and plasmids are listed in table S3. All primers are listed in table S4. *P. syringae* were routinely grown at 27°C on King's medium B (KB) [65]. *E. coli* was routinely grown in lysogeny broth (LB) [66] at 37°C. When appropriate, growth media was supplemented with antibiotics or sugars at the following concentrations: 10 µg/ml tetracycline, 50 µg/ml kanamycin, 25 µg/ml gentamycin, 50 µg/ml rifampicin, nitrofurantoin (NFT) 50 µg/ml, and 5% (w/v) sucrose.

### Deletion and complementation of bacteriocin loci

Colicin-like and R-type-associated syringocin loci were deleted using an overlap-extension approach similar to that described in [67]. Approximately 0.8-1.0 kilobase pairs flanking targeted genes were amplified using *Pfx* polymerase (a proof reading polymerase). Amplified fragments were separated and purified from an agarose gel using the QIAquick Gel Extraction Kit (Qiagen, Valencia, CA) and combined into a subsequent PCR. Combined fragments (in the absence of primers) were cycled 10x, after which BP tailing primer R and F were added and allowed to 20 additional cycles. The desired product was isolated from an agarose gel and combined with the entry vector (pDONR207) in a BP clonase reaction (Invitrogen, Carlsbad, CA). Gentamycin resistant colonies were screened using PCR to confirm the presence of desired DNA fragment. Entry vectors confirmed to harbor the correct fragments were combined with the destination vector (pMTN1907) in an LR clonase reaction (Invitrogen). Following this reaction, kanamycin resistant colonies were screened using PCR, one colony confirmed to contain the correct fragment was used as the mating donor in a tri-parental conjugation with *P. syringae* B728a and DAB44 (mating helper). Matings were plated on KB amended with rifampacin, NFT, and tetracycline. Colonies were screened by PCR to harbor the desired construct. A single confirmed isolate was incubated over night in unamended KB, afterwhich it the resultant culture was serially diluted and spotted onto KB amended with sucrose (counter selection against integrated pMTN1907). Single colonies were picked and confirmed to retain the desired deletion by PCR, as well as to be tetracycline sensitive.

For complementation, genes or operons containing their native promoters were amplified using *Pfx*. Amplified fragments were tailed with BP primers in a subsequent PCR and cloned into pDONR207 in a BP clonase reaction. Isolates confirmed by PCR to contain the desired fragment served as the donor vectors in an LR clonase reaction with either pJC531 (R_reg_) or pBAV226 (R_rbp_) as the destination vector. Plasmid was isolated from one isolate harboring the destination vector confirmed to contain the desired fragment, which was subsequently electroporated into the appropriate *P. syringae* B728a deletion mutant.

### Assay for killing activity

Bacteriocin production was induced by treating log-phase broth cultures with mitomycin C (mitC, 0.5 ng/ml, final concentration), which were allowed to incubate for an additional 12 to 24 hours. Supernatant from induced cultures was collected by centrifuging surviving cells and cell debris and was sterilized either by filtration using a 0.2 µm filter or by adding 5-10 µl of chloroform to culture supernatant. Bacteriocins were occasionally concentrated to increase detectable killing activity using polyethylene glycol (PEG) precipitation. PEG precipitation was performed by adding NaCl (1.0 M final concentration) and PEG 8000 (10% final concentration) to culture supernatants and either incubated in an ice water bath for 1 hour or at 4 °C over night. Samples were centrifuged at 16k G for 30 minutes at 4 °C. Supernatants were decanted and samples were resuspended in 1/10 or 1/100 original volumne in buffer (0.01 M Tris, 0.01 M MgSO_4_, pH 7.0). Residual PEG was removed by extracting twice with equal columes of chloroform. Sterile supernatants (concentrated or not) were tested for killing activity using a standard agar overlay technique [35]. Briefly, 100 µl of log phase broth tester culture was combined with 3 ml molten soft-agar (0.4% agar) which was mixed by vortexing and poured over a standard KB agar plate (1.5% agar). Plates were allowed to solidify for 30 minutes to 1 hour prior to spotting 5 µl of mitC induced, sterilized supernatant. Plates were allowed to incubate for 24-48 hours after which the presence or absence of a clearing zone was determined. To differentiate between bacteriocin-derived clearing and bacteriophage-derived clearing, two tests were performed. First, serial 1/5 dilutions were plated onto the same tester strain. If killing activity was derived from a bacteriophage, dilutions were found where the clearing zone resolved into distinct plaques, whereas killing activity from a bacteriocin did not. Second, material from zones of clearing on a bacterial lawn was transferred using a sterile tooth-pick to a subsequent agar overlay plate of the same tester strain. Clearing zones derived from bacteriophages yielded clearing on the subsequent plate, whereas clearing zones derived from bacteriocins did not yield subsequent clearing.

### Non-R-type bacteriocin prediction and genomic comparison

Colicin-like bacteriocins and their associated immunity proteins (IPs) were predicted by using characterized catalytic domains and IPs (provided by D. Walker, University of Glasgow) as queries using blastp. Additionally, the recently described lectin-like bacteriocin [68] served as a query using blastp. Pathovar genomes were analyzed using both BAGEL2 and BAGEL3 web-based programs [40,41], which predicts a number of different types of bacteriocins, including class I, II, and III bacteriocins. Bacteriocin prediction using BAGEL3 was not performed until after completion of the R-type bacteriocin, thus, the carocin D-like bacteriocin in *Psy* B728a was not targeted for deletion as the source of the killing activity had already been determined.

### R-type bacteriocin prediction and genomic comparison

The genomic region immediately surrounding *trpE* and *trpG* was aligned between *P. syringae* B728a and *P. aeruginosa* PAO1 using progressive Mauve (implemented in Geneious v6.1.8, created by Biomatters, available from http://www.geneious.com/). Additionally, genes were compared between the two strains at the amino acid level using blastp. Hits exhibiting and E-value of 10^-5^ or less were considered significant.

The gene content of the R-type bacteriocins of *P. syringae* B728a and *P. aeruginosa* PAO1 were compared to phage Mu and phage P2, respectively, at the amino acid level using blastp. Hits exhibiting and E-value of 10^-5^ or less were considered significant.

Gene content across the *Pseudomonas* genus was compared using the gene ortholog neighborhood viewer (based on best bidirectional blast hit) at the Integrated Microbial Genomes (IMG) website (https://img.jgi.doe.gov/) with both *trpE* and *trpG* as queries. Comparison of R-type syringacin gene content across *P. syringae* pathovars was also performed using the gene ortholog neighborhood viewer (best bidirectional blast hit). Additionally, genes within the R-type syringacin region that are bacteriophage derived were compared across pathovars using blastp.

### Phylogenetic inference of selected genes

We used a transitive homology approach [47] to identify proteins that exhibited detectable amino acid similarity across the R-type bacteriocin regions from *P. syringage* B728a and *P. aeruginosa* PAO1, as well as from phages Mu and P2. Three proteins were recoverd, including Psyr_4587 (corresponding to gp47 [Mu], gpJ [P2], and PA0618 [*P. aeruginosa*]), Psyr_4589 (corresponding to gp45 [Mu], gpV [P2], and PA0616 [*P. aeruginosa*]), and Psyr_4595 (corresponding to gp39 [Mu], gpFI [P2], and PA0622 [*P. aeruginosa*]). Using blastp searches, homologs representing a range of percent identity with Psyr_4587, Psyr_4589, and Psyr_4595 were collected from the NCBI-nr database, as well from all bacteriophage, *P. syringae*, and *P. aeruginosa* genomes available in the IMG database. Truncated and duplicate (in terms of amino acid identity) sequences were excluded from phylogenetic analyses. Hits were aligned with the MAFFT v1.3.3 plug-in [69] in Geneious v6.1.5 (Biomatters Ltd.) using L-INS-i under Blosum30, with a gap open penalty of 1.53 and an offset value of 0.123.

Maximum-likelihood (ML) heuristic searches were used to estimate phylogenies with RAxML v8.0.9 on the CIPRES Science Gateway under either the WAG + I + G (Psyr_4587, PA0618, gp47, and gpJ), WAG + I + G (Psyr_4589, PA0616, gp45, and gpV), WAG + G + F (Psyr_4595, PA0622, gpL, and gpFI), LG + G + F (Concatenation of multiple sequence alignments for Psyr_4587, Psyr_4589, and Psyr_4595 homologs from *Pseudomonas*), WAG + G (Psyr_4587 and homologs from *Pseudomonas*), LG + G (Psyr_4589 and homologs from *Pseudomonas*), or LG + G + F (Psyr_4595 and homologs from *Pseudomonas*), WAG + I + G + F (gp45 homologs), WAG + G + F (gp47 homologs, or WAG + G + F (L_Psyr homologs), models of evolution as determined by the Bayesian Information Criterion in ProtTest v3.2 [70]. Searches for the phylogenetic reconstruction with the highest likelihood score were performed simultaneously with 1000 rapid bootstrapping replicates. Resulting phylogenies were edited with FigTree v1.3.1 (http://tree.bio.ed.ac.uk/software/figtree/) and Adobe Illustrator.

### Transmission Electron Microscopy

Supernatants from mitomycin C induced cultures of *Psy* B728a, ΔR_reg_, ΔR_struct_ were 10x concentrated using PEG precipitation. Concentrated supernatants were adsorbed onto carbon filter (300 mesh) and stained with uranyl acetate. Samples were visualized using a Phillips CM-12 TEM in the Arizona Health Sciences Center Imaging Core Facility.

## Acknowledgements

This project was supported by by the Agriculture and Food Research Initiative Competitive Grant No. 2015-67012-22773 from the USDA National Institute of Food and Agriculture to K.L.H and by startup funds to D.A.B from the University of Arizona, School of Plant Sciences. T.R. acknowledges support from the National Institutes of Health (K12 GM000708).

We thank Erick Karlsrud, Kevin Dougherty, and Rachel Murillo for their help in generating and testing of various mutants presented in this work. We thank Daniel Walker for supplying bacteriocin catalytic domain sequences. We thank Tony Day at the Arizona Health Sciences Center Imaging Core Facility for assistance with the transmission electron microscopy.

## Supporting Information

**Figure S1.**
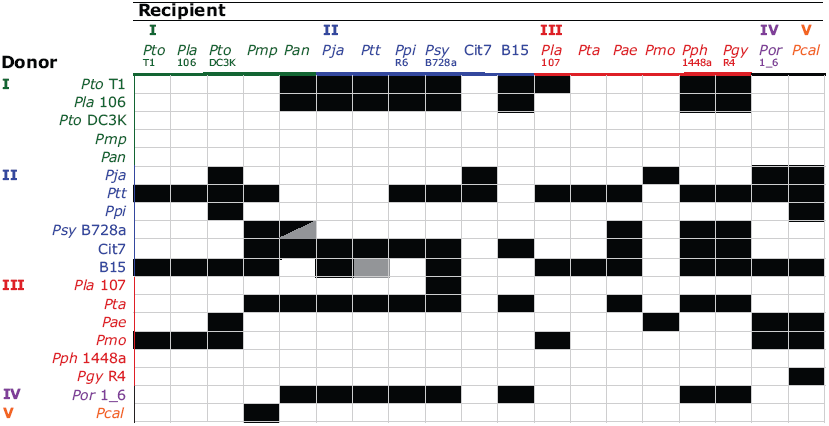
Killing activities of *P. syringae* pathovars. Rows indicate the source of killing activity. Columns target of killing activity. Activities are as follows: no killing activity detected (white box), non-bacteriophage-mediated killing activity (black box), and bacteriophage-mediated killing activity (gray box). Color coded according to clades in **[71]**.

**Figure S2.**
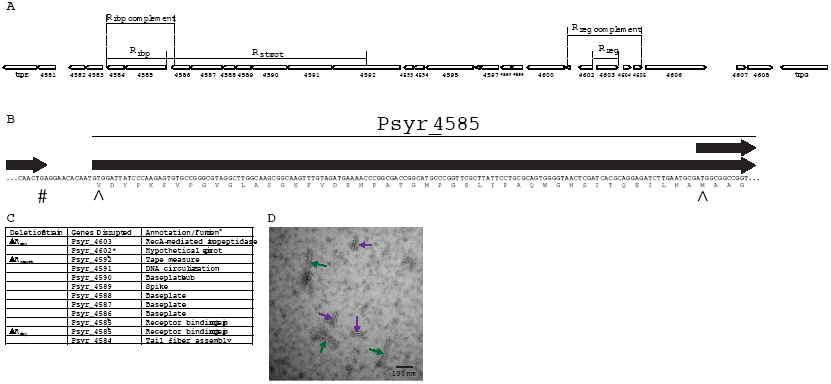
Map of R-type syringacin deletion and complementing constructs. Zones of deletion constructs (ΔR_rbp_, ΔR_struct_, and ΔR_reg_) indicate regions removed in deletion strains. Zones of complement constructs (R_rbp_ complement and R_reg_ complement) indicate regions introduced on replicative plasmids pBAV226 and pMTN41, respectively (A). Original and updated annotated start codon for Psyr_4585. Right most * indicates original start codon (as annotated in **[44]**genome release, GOLD project ID: Gp0000484). Left most * indicates alternative upstream start codon (as annotated in **[72]**genome release, GOLD project ID: Gp004223). Upstream # indicates the stop codon for Psyr_4586 (B). *promoter of gene is removed, coding sequence is unaltered. ^a^ ΔR_reg_ annotations from IMG, ΔR_struct_ and ΔR_rbp_ annotations based on **[48,49]**. ^b^ Prediction based on synteny (C). Transmission electron micrograph (105,000x magnification) mitomycin C induced *Psy* B728a culture supernatant that was concentrated 10x using PEG precipitation (see methods) (D). Green arrows indicate intact tailocins, purple arrows indicate tailocin fragments.

**Figure S3.**
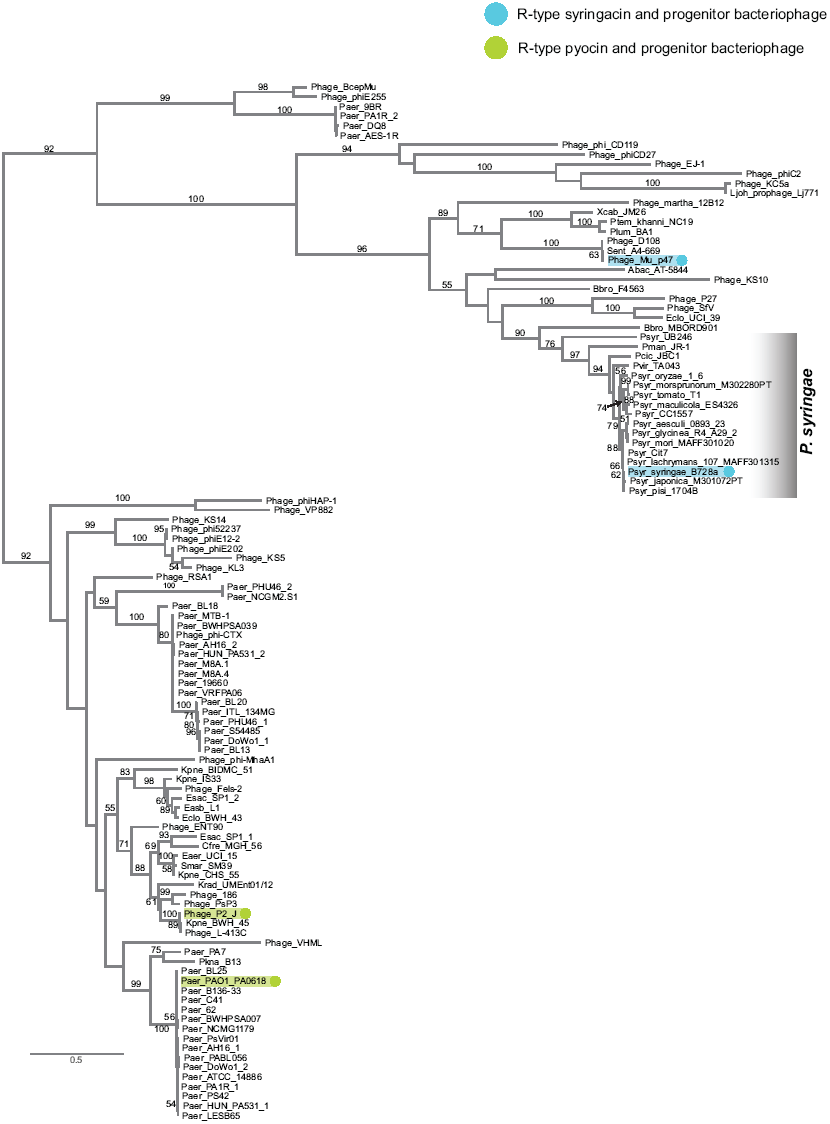
Maximum likelihood phylogeny of Psyr_4587 (*P. syringae* B728a), PA0618 (*P. aeruginosa* PAO1), gp47 (Mu), and gpJ (P2) and related protein sequences from bacteria and bacteriophages. Teal highlighted nodes indicate Psyr_4587 and gp47, whereas green highlighted nodes indicate PA0618 and gpJ. Values on branches indicate bootstrap support (1000 bootstrap replicates) for those clades (only values of 50 or greater are shown). A monophyletic clade including all *P. syringae* orthologs is indicated. See table S2 for accessions or IMG GeneIDs associated with each sequence.

**Figure S4.**
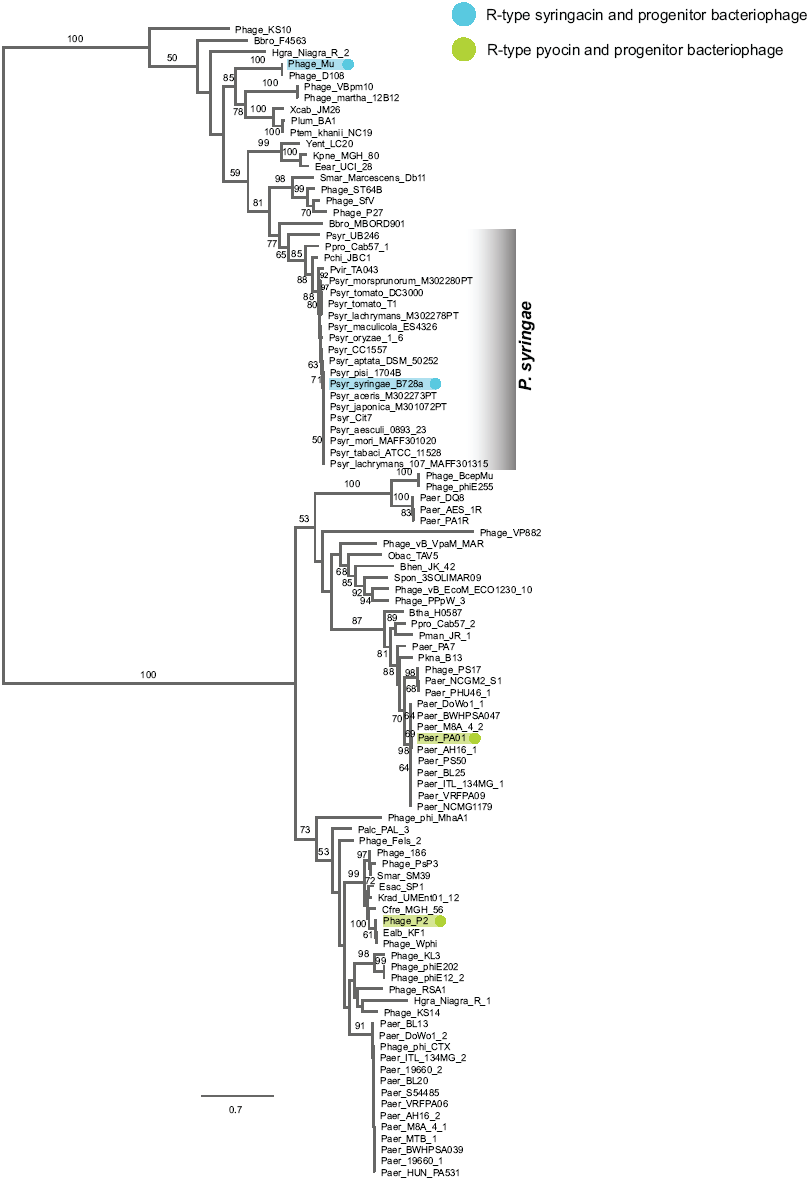

Maximum likelihood phylogeny of Psyr_4595 (*P. syringae* B728a), PA0622 (*P. aeruginosa* PAO1), gpL (Mu), and gpFI (P2) and related protein sequences from bacteria and bacteriophages. Teal highlighted nodes indicate Psyr_4595 and gpL, whereas green highlighted nodes indicate PA0622 and gpFI. Values on branches indicate bootstrap support (1000 bootstrap replicates) for those clades (only values of 50 or greater are shown). A monophyletic clade including all *P. syringae* orthologs is indicated. See table S2 for accessions or IMG GeneIDs associated with each sequence.

**Figure S5.**
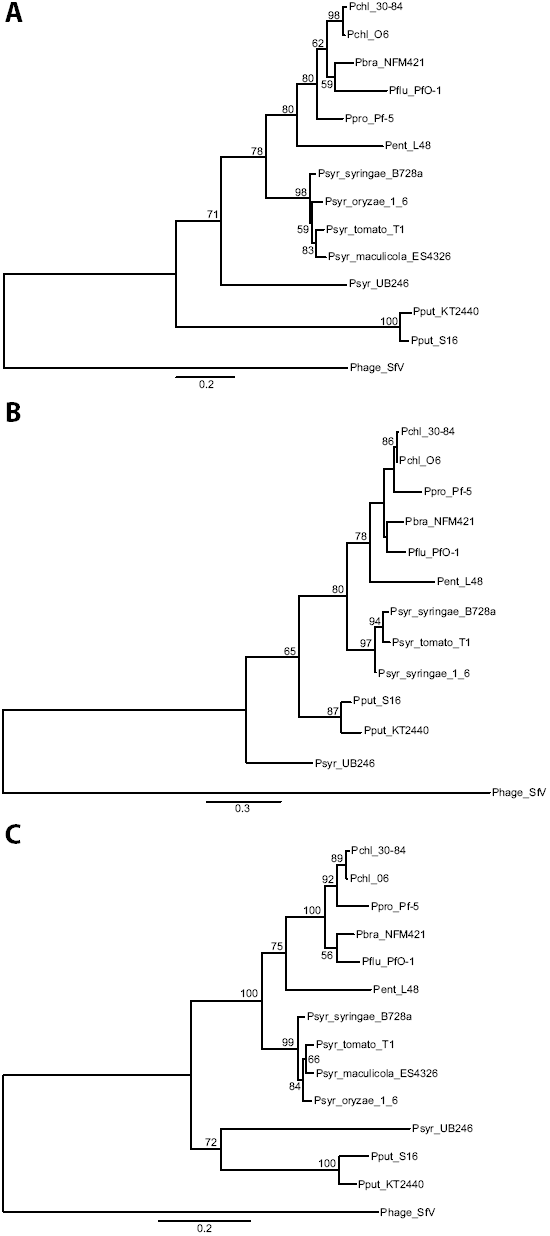
Maximum likelihood phylogeny of amino acid sequence of Psyr_4587 (A), Psyr_4589 (B), Psyr_4595 (C) and their homologs from *Pseudomonas* strains harboring an R-type syringocin-like region. Values indicate bootstrap support (out of 1000 replicates, only values greater than 50 are shown). The following strains are Pchl 30-84 (*P. chlororaphis*30-84), Pchl O6 (*P. chlororaphis* O6), Pbra NFM421 (*P. brasicacearum* NFM421), Pflu PfO-1 (*P. fluorescens* PfO-1), Ppro Pf-5 (*P. protegens* Pf-5), Pent L48 (*P. entomophila* L48), Psyr B728a (*P. syringae* pv. syringae B728a), Psyr 1_6 (*P. syringae* pv. oryzae 1_6), Psyr T1 (*P. syringae* pv. tomato T1), Pcan ES4326 (*P. cannabina* pv. alisalensis ES4326), Psyr UB246 (*P. syringae* UB246), Pput KT2440 (*P. putida* KT2440), Pput S16 (*P. putida* S16). Phage SfV (bacteriophage SfV, outgroup).

**Table S1.**
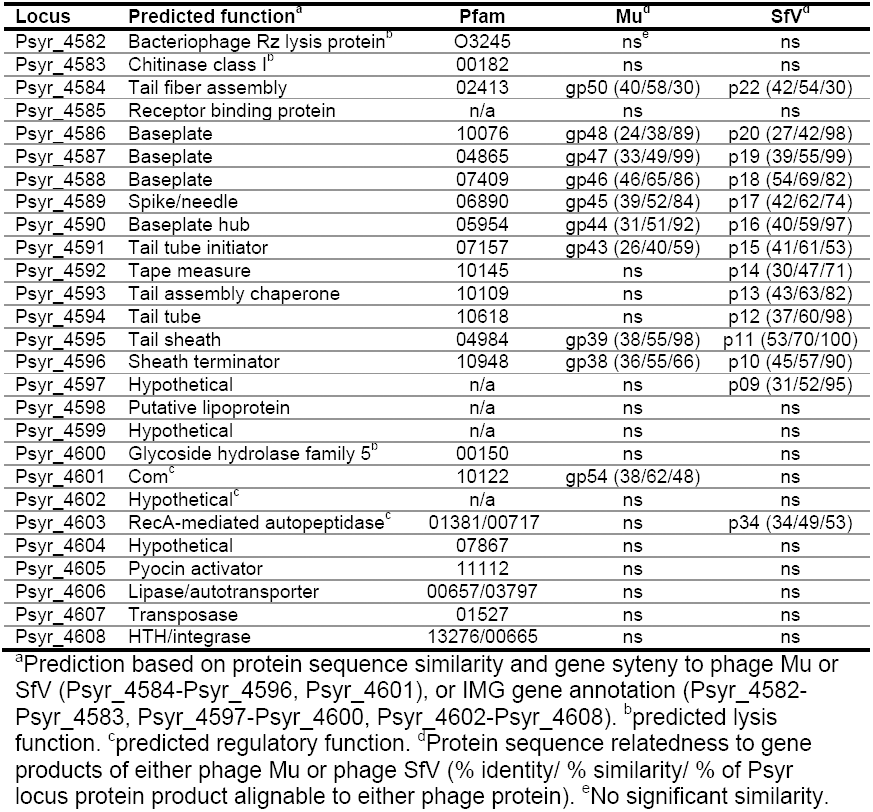
Gene content of the R-syringacin region from *Psy* B728a.

Table S2. Sequence designations used in figures 6, S3, and S4.

**Table S3.** Strains and Plasmids.

| Strain or plasmid | Description or Relevant Characteristics | Antibiotics | Reference |
| --- | --- | --- | --- |
| *E. coli* strains |  |  |  |
| TOP10 | F- *mcrA* $\Delta$(*mrr-hsdRMS-mcrBC*) ϕ80*lacZ*ΔM15 Δ*lacX74* *recA1* *araD139* $\Delta$(*araleu*) 7697 *galU* *galK* *rpsL* (Str<sup>R</sup>) *endA1* *nupG* | | Invitrogen |
| DAB44 | *E. coli* harboring pRK2013, (conjugation helper for triparental mating) |  | [66] |
| *P. syringae* strains |  |  |  |
| *Pto* T1 | *P. syringae* pv. tomato T1 |  | [67] |
| *Pla* 106 | *P. syringae* pv. lachrymans MAFF302278PT |  | [63] |
| *Pto* DC3K | *P. syringae* pv. tomato DC3000 |  | [68] |
| *Pmp* | *P. syringae* pv. morsprunorum MAFF302280PT |  | [63] |
| *Pan* | *P. syringae* pv. actinidiae MAFF302091 |  | [63] |
| *Pja* | *P. syringae* pv. japonica MAFF 301072 PT |  | [63] |
| *Ptt* | *P. syringae* pv. aptata DSM50252 |  | [63] |
| *Ppi* | *P. syringae* pv. pisi 1704B |  | [63] |
| *Psy* B728a | *P. syringae* pv. syringae B728a |  | [44] |
| *Pac* | *P. syringae* pv. aceris MAFF302273PT |  | [63] |
| *Cit7* | *P. syringae* *Cit7* |  | [63] |
| *Pla* 107 | *P. syringae* pv. lachrymans MAFF301315 |  | [63] |
| *Pta* | *P. syringae* pv. tabaci ATCC11528 |  | [63] |
| *Pae* | *P. syringae* pv. aesculi 0893_23 |  | [63] |
| *Pmo* | *P. syringae* pv. mori MAFF301020 |  | [63] |
| *Pph* 1448A | *P. syringae* pv. phaseolicola 1448A |  | [69] |
| *Pgy* R4 | *P. syringae* pv. glycinea A29-2 |  | [63] |
| *Por* 1_6 | *P. syringae* pv. oryzae |  | [63] |
| *Pcal\** | *P. cannabina* pv. alisalensis |  | [63] |
| Δ0310 | *Psy* B728a harboring a markerless deletion of *Psy*<sub>r</sub>_0310 and *Psy*<sub>r</sub>_0309 | | This study |
| Δ4651 | *Psy* B728a harboring a markerless deletion of *Psy*<sub>r</sub>_4651 and *Psy*<sub>r</sub>_4652 | | This study |
| ΔS-type | *Psy* B728a harboring markerless deletions of *Psy*<sub>r</sub>_0310, *Psy*<sub>r</sub>_0309, *Psy*<sub>r</sub>_4651, and *Psy*<sub>r</sub>_4652 | | This study |
| ΔR<sub>reg</sub> | *Psy* B728a harboring a markerless deletion of *Psy*<sub>r</sub>_4603 and the promoter of *Psy*<sub>r</sub>_4602 (Fig. S1) | | This study |
| ΔR<sub>struct</sub> | *Psy* B728a harboring a markerless deletion of *Psy*<sub>r</sub>_4586 through *Psy*<sub>r</sub>_4592 and the promoter of *Psy*<sub>r</sub>_4585 (Fig. S1) | | This study |
| ΔR<sub>rbp</sub> | *Psy* B728a harboring a markerless deletion of *Psy*<sub>r</sub>_4585 and *Psy*<sub>r</sub>_4584 (Fig. S1) | | This study |
| ΔR<sub>reg</sub> + pR<sub>reg</sub> | ΔR<sub>reg</sub> harboring pR<sub>reg</sub> complementation vector (Kan<sup>r</sup>, Fig. S1) | | This study |
| ΔR<sub>rbp</sub> + pR<sub>rbp</sub> | ΔR<sub>rbp</sub> harboring pR<sub>rbp</sub> complementation vector (Tet<sup>r</sup>, Fig. S1) | | This study |
| Plasmids |  |  |  |
| pJC531 | Replicative destination vector harboring *attR* cloning sites | Kan<sup>R</sup> | [70] |
| pMTN1907 | Non-replicative destination vector harboring *attR* cloning sites | Kan<sup>R</sup>, Tet<sup>R</sup>, Cam<sup>R</sup>, Suc<sup>S</sup> | [71] |
| pDONR207 | Entry vector harboring *attP* cloning sites | Gent<sup>R</sup>, Cam<sup>R</sup> | Invitrogen |
| pBAV226 | Replicative destination vector with constitutive nptII promoter | Tet<sup>R</sup> | [72] |
| pR<sub>reg</sub> | *Psy*<sub>r</sub>_4603 and *Psy*<sub>r</sub>_4602 plus flanking sequence in pJC531 (Fig. S1) |  | This study |
| pR<sub>rbp</sub> | *Psy*<sub>r</sub>_4584 and *Psy*<sub>r</sub>_4585 complementation construct (including upstream alternative start codon) in pBAV226 (Figs. S1 and S2) |  | This study |
| pKLHECOEN3 | *Psy*<sub>r</sub>_4585 and *Psy*<sub>r</sub>_4584 deletion construct integrated into pDONR207 |  | This study |
| pKLHECODE5 | *Psy*<sub>r</sub>_4585 and *Psy*<sub>r</sub>_4584 deletion construct integrated into pMTN1907 |  | This study |
| pKLHECOEN11 | R<sub>rbp</sub> complementation construct integrated into pDONR207 |  | This study |
| pKLHECOEN14 | *Psy*<sub>r</sub>_0310 and *Psy*<sub>r</sub>_0309 deletion construct integrated into pDONR207 |  | This study |
| pKLHECODE15 | *Psy*<sub>r</sub>_0301 and *Psy*<sub>r</sub>_0309 deletion construct integrated into pMTN1907 |  | This study |
| pKLHECOEN16 | *Psy*<sub>r</sub>_4651 and *Psy*<sub>r</sub>_4652 deletion construct integrated into pDONR207 |  | This study |
| pKLHECODE17 | *Psy*<sub>r</sub>_4651 and *Psy*<sub>r</sub>_4652 deletion construct integrated into pMTN1907 |  | This study |
| pKLHECOEN18 | R<sub>struct</sub> deletion construct integrated into pDONR207 |  | This study |
| pKLHECODE19 | R<sub>struct</sub> deletion construct integrated into pMTN1907 |  | This study |
| pKLHECOEN20 | R<sub>reg</sub> deletion construct integrated into pDONR207 |  | This study |
| pKLHECODE21 | R<sub>reg</sub> deletion construct integrated into pMTN1907 |  | This study |
| pKLHECOEN22 | R<sub>reg</sub> complementation construct integrated into pDONR207 |  | This study |

\*Also known as *P. syringae* pv. *maculicola* ES4326

**Table S4.**
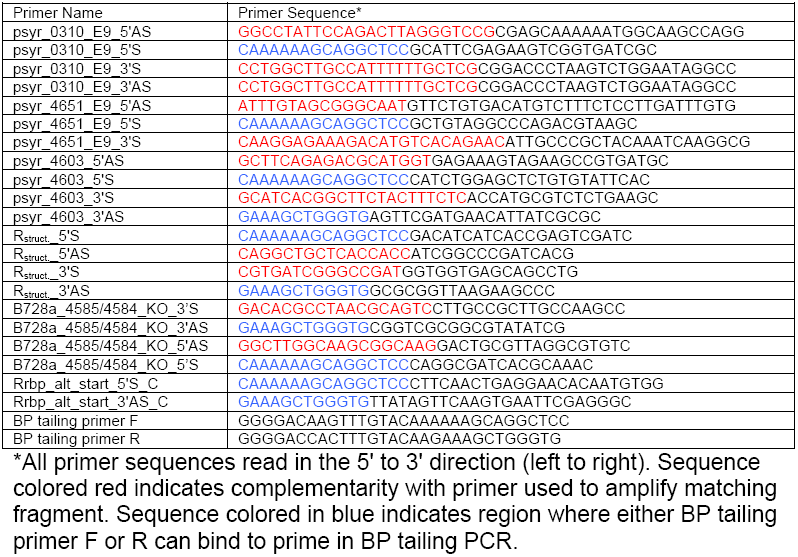
Primers.

